# Natural and planted forests differ in the mechanisms regulating productivity and stability

**DOI:** 10.64898/2026.09.10.750640

**Authors:** Mingjie Chen, Daoli Peng, Elia Vangi, Mauro Morichetti, Alessio Collalti

## Abstract

Forest productivity and stability underpin ecosystem functioning and services, yet whether the drivers and mechanisms regulating these properties differ between natural and planted forests at large spatial scales remains unclear. Using data from 2,389 forest plots across China and a novel analytical framework, we systematically compared productivity and stability between natural and planted forests and identified their dominant drivers and underlying mechanisms across 38 biotic, structural, climatic, and edaphic variables. We found that natural forests were 14% more productive and 34% more stable than planted forests, although these differences varied across ecoregions and stand development stages. Productivity in natural forests was primarily driven by functional diversity and functional traits, supporting the niche complementarity and mass-ratio hypotheses, whereas nitrogen availability emerged as the dominant regulator of productivity in planted forests. Likewise, stability shifted from diversity-related regulation in natural forests to stronger abiotic control in planted forests, while functional diversity remained a key stabilizing mechanism in both forest types, supporting the insurance hypothesis. We also found that planted forests were more sensitive to hydroclimatic variability, indicating greater vulnerability to drought. Our findings reveal a fundamental shift in the mechanisms regulating forest productivity and stability from diversity-mediated control in natural forests to stronger abiotic regulation in planted forests, highlighting the importance of integrating biodiversity conservation and nitrogen management to sustain forest productivity and stability under future climate change.

## 1. Introduction

Forest ecosystems constitute the most important reservoir of terrestrial biodiversity and represent one of the largest carbon pools on earth, storing more than 80% of aboveground vegetation carbon and over 70% of global soil organic carbon (Six et al. 2002; Pan et al. 2011, 2024). Through carbon sequestration, climate regulation, nutrient cycling, and habitat provision, forests play a central role in sustaining ecosystem functioning and human well-being (Bonan 2008). However, ongoing deforestation, forest degradation, and climate change are increasingly threatening forest ecosystems worldwide, leading to substantial biodiversity loss and raising concerns about their capacity to maintain ecosystem functioning under future environmental change (Feng et al. 2016; Yu et al. 2019; Hills et al. 2023). In this context, identifying the determinants of forest productivity and stability, which represent two fundamental dimensions of ecosystem functioning, is essential for developing sustainable forest management strategies and mitigating the impacts of climate change on forest ecosystems (Vangi et al. 2024a).

Forest productivity reflects the rate at which vegetation assimilates carbon from the atmosphere and partially converts it into organic matter (Friedlingstein et al. 2023), whereas stability describes the ability of ecosystems to maintain functioning in the face of environmental variability and disturbance (Ding et al. 2023). Both properties are regulated by a variety of biotic and abiotic factors. Among biotic drivers, tree diversity has long been recognized as a key determinant (Mori et al. 2017; Homeier & Leuschner 2021). Forests with higher tree species diversity often exhibit greater productivity and temporal stability owing to niche complementarity and species asynchrony, although the strength and direction of these relationships can vary across spatial and temporal scales (Liang et al. 2016; Mori et al. 2017; Bongers et al. 2021). Beyond species diversity, functional composition and functional diversity influence forest functioning through trait-mediated resource acquisition strategies and mass-ratio effects (Grime 1998; Yuan et al. 2018). Increasing evidence also highlights the importance of structural diversity, which can enhance canopy stratification, resource partitioning, and stand-level functioning across tropical, subtropical, and temperate forests (Dieler et al. 2017; Schnabel et al. 2019; Ouyang et al. 2023). Stand attributes such as biomass, density, and age further shape these two properties by altering resource availability and competitive interactions between individuals (Vangi et al. 2024a, b). Abiotic factors including climate, soil properties, and topography regulate forest functioning by constraining resource availability and growth conditions (Bardgett & Van Der Putten 2014; Michaletz et al. 2014; García-Palacios et al. 2018). Moreover, global change drivers such as nitrogen deposition, elevated atmospheric CO_2_ concentrations, and increasing drought frequency are exerting increasingly strong influences on forest productivity and stability worldwide (Bobbink et al. 2010; Anderegg et al. 2020; Du et al. 2020).

Although the drivers of forest productivity and stability have been extensively investigated, the role of forest origin has received comparatively little attention. Forest origin fundamentally shapes community assembly, species composition, and stand structure, potentially altering the ecological processes that regulate ecosystem functioning. Natural forests develop through succession and natural regeneration, typically resulting in greater taxonomic and structural complexity (Hua et al. 2022; Pei et al. 2026), whereas planted forests are established through human intervention and are often characterized by simplified species composition and more homogeneous stand structure (Barlow et al. 2007; Brockerhoff et al. 2008). These differences may lead to contrasting ecosystem functioning and responses to environmental and climate change. For example, natural forests often exhibit greater carbon storage capacity, higher water-use efficiency, and stronger resistance to drought stress than planted forests (Yu et al. 2019; Zhong et al. 2021; Sun et al. 2024). However, productivity and stability have rarely been evaluated simultaneously across forest origins, and it remains unclear whether the drivers and mechanisms regulating these two fundamental properties differ between natural and planted forests at large spatial scale. China provides an ideal study system for addressing these questions. Since the 1970s, large-scale ecological restoration and afforestation programs, including the Three-North Shelterbelt Program, the Grain for Green Program, and nationwide land-greening initiatives, have substantially expanded planted forests across the country. As a result, China now hosts the largest area of planted forests in the world, accounting for approximately 27% of the global total (80 million ha) (FAO and UNEP, 2020). In parallel, natural forests remain widespread owing to long-term conservation efforts, including the Natural Forest Conservation Program, and currently account for approximately 64% of China’s forest area, corresponding to about 138.7 million ha (National Forestry and Grassland Administration, 2019).

Based on the fundamental ecological differences between natural and planted forests, and on previous theoretical and empirical evidence, we hypothesized that: (1) natural forests generally exhibit higher productivity and stability than planted forests across environmental gradients; and (2) productivity and stability in natural forests are primarily regulated by biotic factors such as tree diversity and stand structure, whereas abiotic factors, including nitrogen deposition and soil nutrient availability, exert stronger influences in planted forests. To test these hypotheses, we compiled a nationwide dataset comprising 2,389 forest plots spanning natural and planted forests across China and characterized by 38 biotic and abiotic variables. We developed a novel analytical framework that combines the XGBoost machine learning algorithm, Shapley Additive Explanations (SHAP) analysis, and structural causal model (SCM) to identify key driving factors and to infer causal mechanisms regulating forest productivity and stability. By explicitly comparing the determinants of productivity and stability between natural and planted forests, our study reveals how forest origin mediates productivity and stability and provide insights for forest restoration and sustainable forest management under future global change.

## 2. Materials and Methods

### 2.1. Forest inventory data

We compiled a nationwide forest inventory dataset comprising 2,389 sample plots (20 m × 30 m), all based on ground-based measurements conducted between 2013 and 2019. The dataset integrates plots obtained from the National Earth System Science Data Center, the National Science & Technology Infrastructure of China (http://www.geodata.cn) with those collected through our own field surveys. The plots span all major forest climatic zones across mainland China, excluding Hong Kong, Taiwan, and Macau (Figure 1). Natural forest plots subject to natural or anthropogenic disturbances were excluded, resulting in a final total of 1,273 natural and 1,116 planted plots. Within each plot, all trees with a diameter at breast height (DBH) ≥ 5 cm were surveyed for species identity, DBH, tree height, and age class. All measurements followed standardized forest inventory protocols (Chinese National Standard, GB/T 30363–2013; Chinese Forestry Standard, LY/T 3128–2019). These measurements were used to derive stand-level structural attributes, including stand density, mean DBH, and mean tree height. Plot-level total biomass was estimated as the sum of aboveground and belowground components using species-specific allometric equations, following the national standard “Tree biomass models and related parameters to carbon accounting for major tree species (GB/T 43648–2024)”.

**FIGURE 1.**
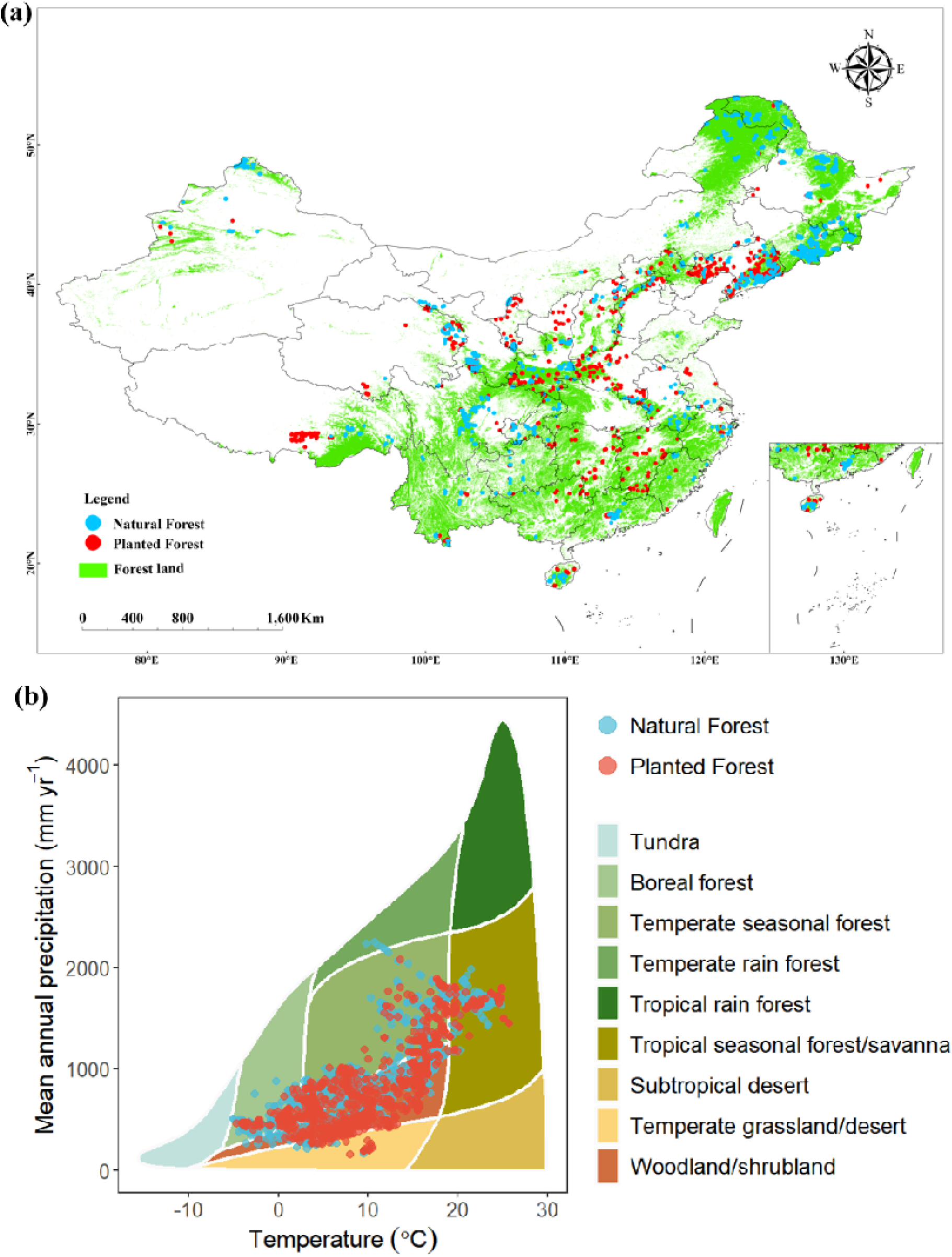
(a) Spatial distribution of natural forest and planted forest plots across China used in this study. Blue circles represent natural forest plots, red circles indicate planted forest plots, and green areas delineate forest land coverage across the country. (b) Whittaker biome diagram illustrating the climatic space occupied by forest plots in the database, plotted as a function of mean annual temperature (°C, x-axis) and mean annual precipitation (cm, y-axis). Each point represents an individual forest plot, colored according to elevation (m a.s.l.), ranging from low-elevation sites (red, ∼0 m) to high-elevation sites (green, >4000 m). Background polygons delineate the nine major Whittaker biomes.

### 2.2. Forest productivity and stability data

This study employs the kernel normalized difference vegetation index (kNDVI) as a proxy for forest productivity (Forzieri et al. 2022; Xi et al. 2024; Pickering et al. 2025). kNDVI has been shown to be strongly correlated with gross and net primary productivity and to outperform traditional vegetation indices by reducing saturation effects and noise sensitivity across broad environmental gradients and spatiotemporal scales (e.g., normalized difference vegetation index (NDVI), enhanced vegetation index (EVI), and near-infrared reflectance of vegetation (NIRv)) (Camps-Valls et al. 2021; Qi et al. 2023). The kNDVI was calculated as follows:

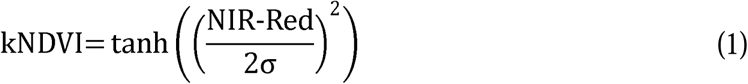

Where NIR and Red represent the reflectance of the near-infrared and red bands, respectively, and σ is a length-scale parameter that determines the sensitivity of the index to variations in vegetation density. σ was specified as 0.5 × (NIR + Red) (Camps-Valls et al. 2021).

All available Landsat 5, 7, and 8 surface reflectance imagery covering mainland China from 2001 to 2020 was acquired via the Google Earth Engine (GEE) platform. Standard preprocessing procedures were applied, including atmospheric correction and the removal of cloud, shadow, snow, and rainfall contamination using quality assurance masks (Foga et al. 2017). To minimize the adverse effects of clouds, aerosols, and shadows, monthly maximum-value composite kNDVI images were first generated, and annual kNDVI was then calculated as the mean of the monthly composites for each year (Holben 1986; Xu et al. 2024; Yu et al. 2025). A circular buffer with a radius of 15 m was applied around the geographic center of each forest plot to match the plot size (667 m²), and mean annual kNDVI values within the buffer were extracted for each year from 2001 to 2020.

Forest stability was quantified as the temporal stability of kNDVI, calculated as the ratio of the mean annual kNDVI to its interannual variability over the study period (2001–2020):

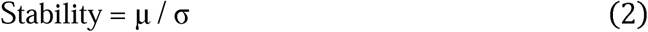

where μ represents the mean annual kNDVI and σ denotes its standard deviation. Because long-term directional trends in vegetation productivity can bias estimates of stability (Huang & Xia 2019; Ren et al. 2023), we applied a detrending procedure before calculating stability. Specifically, a linear regression was fitted between annual kNDVI and year for each plot, and the standard deviation of the regression residuals was used to replace σ in the stability calculation. This approach isolates interannual variability from long-term trends, focusing on fluctuations around the mean state rather than total variability.

### 2.3. Tree diversity indices calculation

In this study, forest biodiversity was characterized using three complementary dimensions: tree species diversity, functional diversity, and structural diversity. Tree species diversity was quantified as species richness, defined as the total number of tree species within each plot. Functional diversity was calculated based on four key plant functional traits closely linked to forest productivity and stability: specific leaf area (SLA, mm^2^ mg^-1^), leaf area (LA, mm^2^), leaf nitrogen content (LN, mg g^-1^), and leaf dry matter content (LDMC, g g^-1^) (Garnier et al. 2004; Wright et al. 2004). Trait data were obtained from the TRY plant trait database (Kattge et al. 2020) and the China plant trait database version 2 (Wang et al. 2022) according to tree species recorded in each plot. When trait values were missing for certain species, they were imputed using the R package funspace (Carmona et al. 2024), which estimates trait values based on multivariate trait relationships. Functional diversity was quantified using functional dispersion (FDis), which represents the mean distance of individual species to the centroid of all species in a multidimensional trait space and is independent of species richness, making it suitable for both species-rich natural forests and species-poor plantation forests (Laliberté & Legendre 2010; Yuan et al. 2020). In addition, community-weighted mean (CWM) values for each trait were calculated with species weighted by their relative basal area within each plot. Structural diversity was quantified using the Gini coefficient of individual tree basal area within each plot (Lexerød & Eid 2006; Duduman 2011; Padilla-Martínez et al. 2024).

### 2.4. Environmental data

To comprehensively assess the abiotic controls on forest productivity and stability in natural and planted forests, we compiled a set of 24 abiotic variables representing climate, global change drivers, drought conditions, topography, and soil properties (Table S1), in addition to the biotic factors described above. Climatic variables, including annual mean temperature (MAT) and annual mean precipitation (MAP), were derived from the 1 km monthly temperature and precipitation dataset for China (Peng et al. 2019) for the period 2001–2020, and aggregated to annual means.

Drought conditions were represented by the Standardized Precipitation Evapotranspiration Index (SPEI), obtained from the GPRChinaSPEI1km dataset (He et al. 2025). We used 3-month time-scale SPEI values to capture short-term moisture stress relevant to vegetation growth and calculated plot-level means for 2001–2020. Atmospheric nitrogen deposition data were derived from a gridded dataset at 0.25° × 0.25° spatial resolution (Zhou et al. 2023), and mean values for 2005–2020 were extracted for each plot.

Atmospheric CO concentration data were obtained from a 1-km gridded dataset (He et al. 2023), and plot-level averages were calculated for 2015–2020. Topographic variables, including elevation, slope, and aspect, were derived from the NASADEM product (NASA JPL, 2020) at 30 m resolution using the GEE platform.

Soil moisture was obtained from the GLASS SM product at 1 km spatial resolution (Zhang et al. 2023), and mean values for 2001–2020 were calculated. Soil physicochemical properties were extracted from the China dataset of soil properties for land surface modelling version 2 (CSDLv2) (Shi et al. 2025) at 1 km resolution, including sand, silt, clay, rock fragment content, porosity, bulk density, pH, soil organic carbon, cation exchange capacity, total nitrogen, total phosphorus, total potassium, alkali-hydrolysable nitrogen, available phosphorus, and available potassium. All soil variables represent average conditions for the 0–200 cm soil profile.

### 2.5. Statistical analyses

The Analysis of variance (ANOVA) was first used to quantify differences in forest productivity and stability between natural and planted forests across ecological zones and forest age groups.

To identify the key drivers of forest productivity and stability in natural and planted forests, we employed the Extreme Gradient Boosting (XGBoost) machine learning algorithm, combined with SHAP analysis for model interpretation. XGBoost is a scalable tree-based ensemble method that effectively captures complex nonlinear relationships and interactions among predictors while minimizing overfitting and noise sensitivity (Chen & Guestrin, 2016). SHAP, grounded in cooperative game theory (Lundberg & Lee, 2017), provides a unified framework for interpreting black-box models by quantifying the marginal contribution of each predictor to model predictions and revealing their functional relationships with response variables. Before model construction, multicollinearity among the 38 biotic and abiotic predictors was assessed using the variance inflation factor (VIF). Variables with VIF > 5 were excluded to reduce collinearity effects. XGBoost models were then trained on the filtered predictor set, and SHAP analysis was applied to quantify variable contribution using mean absolute SHAP values. Hyperparameters were tuned via GridSearchCV method and the optimal parameters were provided in Table S2. Marginal effect plots were generated to examine the direction and strength of relationships between individual predictors and forest productivity or stability. Positive SHAP values indicate positive contributions to model predictions, whereas negative values indicate negative contributions. All these analyses were conducted using the *scikit-learn* and *shap* libraries in Python (version 3.11.6).

Because XGBoost and SHAP analyses are inherently predictive and do not establish causal relationships, we further implemented a causal inference framework based on SCM (Pearl 2009), to disentangle the mechanisms underlying forest productivity and stability in natural and planted forests. Causal structures were specified using directed acyclic graphs (DAGs), which explicitly represent hypothesized causal relationships in a given system. The validity of each DAG was evaluated using a model–data consistency test, with a pass threshold of 0.3 as in previous studies (Chen et al. 2024; McLean et al. 2025). The DAGs imply a set of independencies and conditional independencies among variables, which should be supported by the data if the DAG adequately represents the underlying data-generating process. These implied independencies were identified and statistically tested using the *dagitty* package in R, based on tests of zero (partial) correlation (Ankan et al. 2021). A DAG was considered consistent with the data when none of the implied independencies were statistically refuted. Otherwise, the DAG was iteratively revised to account for missing or incorrect causal pathways (Arif & MacNeil 2023). The final DAGs that passed the consistency test are presented in Figure S1. Based on the validated DAGs, minimal adjustment sets were identified using the backdoor criterion. The backdoor criterion identifies sets of variables that block all noncausal paths between exposure and outcome while preserving causal pathways (Arif & MacNeil 2023). When multiple valid adjustment sets were available, selection was guided by data availability and measurement quality.

Finally, generalized additive mixed models (GAMMs) were fitted to estimate the causal effects of the most influential predictors identified by the XGBoost–SHAP analysis. The structure of each regression model was determined based on the minimal adjustment sets. Prior to GAMMs fitting, spatial autocorrelation in model residuals was assessed using Moran’s I test. Significant spatial autocorrelation effect was detected, and empirical semivariograms were used to determine the distance at which spatial autocorrelation diminished. The resulting thresholds were approximately 2 km for productivity models and 1.2 km for stability models in both natural and planted forests. Accordingly, plot-level group identifiers (groupID) were constructed from these distances and incorporated as random effects in the GAMMs to account for spatial autocorrelation (Aguirre Gutiérrez et al. 2022; Ding et al. 2024). In addition, ecoregions were included as random effects to control for the potential non-independence among plots within the same ecoregion. All analyses were conducted in R using the *dagitty* (Ankan et al. 2021), *mgcv* (Wood 2011), *sf* (Pebesma 2018), *dplyr* (Wickham et al. 2026), *ape* (Paradis & Schliep 2019), and *gstat* (Gräler et al. 2016) packages.

## 3. Results

### 3.1 Forest productivity and stability in natural forest and planted forest across China

The ANOVA analysis showed significant differences in productivity and stability between forest types at the national scale (p < 0.001), with natural forests exhibiting higher productivity (0.56 ± 0.08 vs. 0.49 ± 0.14) and stability (14.48 ± 9.03 vs. 10.84 ± 7.60) than planted forests (Figure 2a,b). A positive nonlinear relationship between forest productivity and stability was observed in both natural and planted forests (Figure 2c). The stability increased gradually at low productivity levels and more rapidly at higher productivity levels. Interestingly, in natural forests, the stability tended to be slightly higher than that in planted forests at both low and high productivity levels, whereas the two forest types showed similar patterns at intermediate productivity levels, with substantial overlap between them.

**FIGURE 2.**
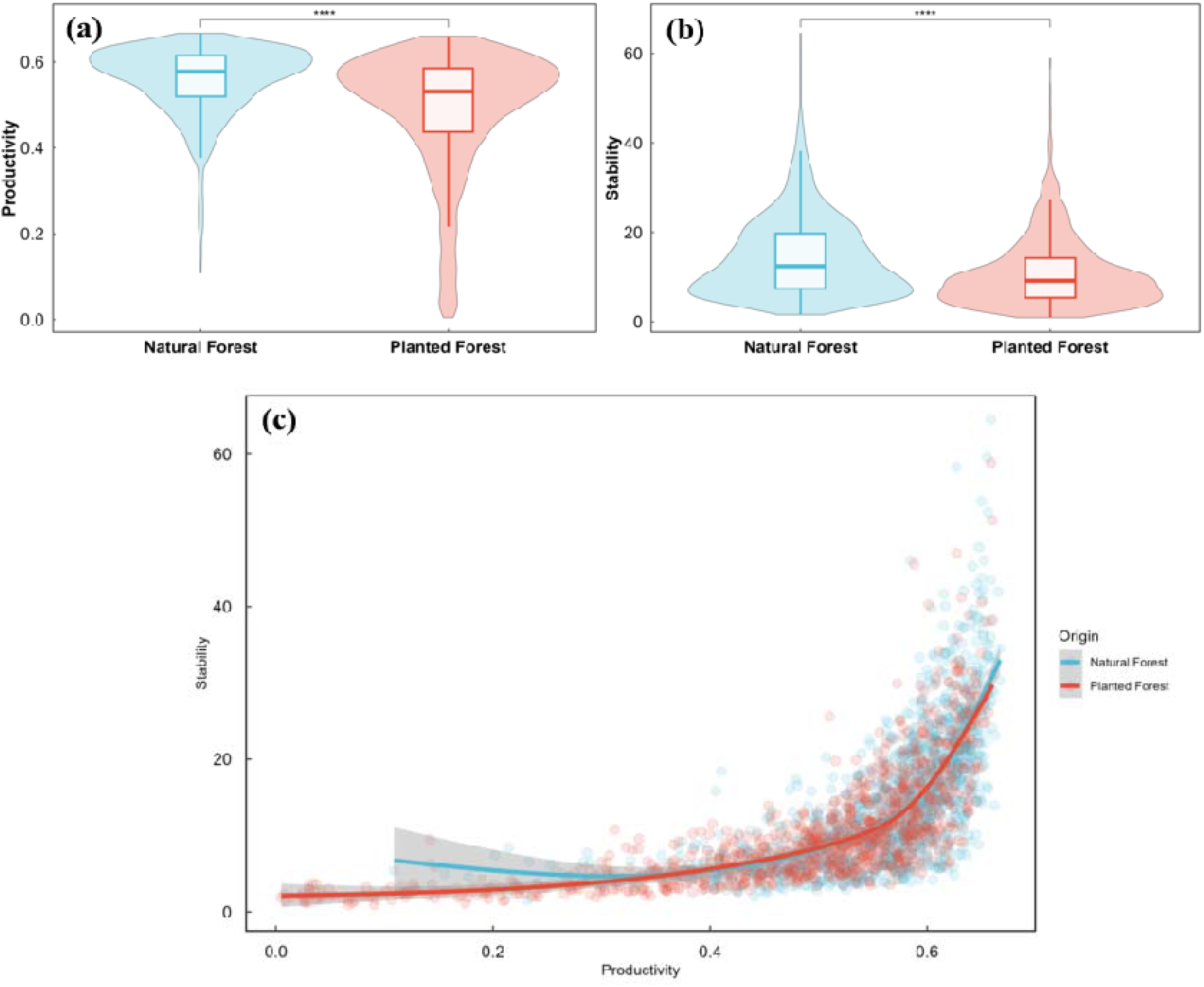
Differences in productivity and stability between natural and planted forests across China. Violin plots show the distributions of (a) productivity and (b) stability in natural forests (blue) and planted forests (red). Embedded boxplots indicate the median, interquartile range, and data range. Symbols denote significant differences between forest types (****, p < 0.001). (c) Relationships between productivity and ecosystem stability in natural forests (blue) and planted forests (red). Points represent individual forest plots, solid lines represent GAM fits, and shaded areas indicate 95% confidence intervals.

The magnitude and direction of these differences varied among ecoregions (Figure S3). Natural forests generally exhibited higher productivity in plateau, temperate, and subtropical coniferous-dominated ecoregions, whereas productivity differences were negligible elsewhere. Patterns of stability showed stronger regional heterogeneity. Natural forests were more stable than planted forests across most subtropical and broadleaf-dominated ecoregions, while planted forests exceeded natural forests only in the Middle Temperate Coniferous Forest.

Forest age further influenced the differences in forest productivity and stability between natural and planted forests (Figure S4). Natural forest productivity was significantly higher than that of planted forests in young, middle-aged, near-mature, and mature stands, but the difference became non-significant in overmature stands. Similarly, stability was significantly higher in natural forests than in planted forests at the young, middle-aged, and near-mature stages, whereas no significant differences were observed at the mature and overmature stages.

### 3.2 Drivers of forest productivity and stability in natural forest and planted forest

To identify the dominant drivers of forest productivity and stability between natural and planted forests, we evaluated the relative importance of 38 predictors representing tree diversity, stand structure, functional traits, climate, soil and topography. In natural forests (Figure 3a), productivity was associated with tree diversity (30.1%), soil properties (20.9%), and functional traits (20.7%). At the individual-predictor level, functional diversity (FD) and the community-weighted mean specific leaf area (CWM SLA) emerged as the dominant drivers, followed by soil sand content, species richness (SR), and nitrogen deposition. In contrast, productivity in planted forests (Figure 3c) was more strongly associated with abiotic factors, with soil properties accounting for 41.3% and climatic factors for 25.1%. Among single predictors, soil alkali-hydrolyzable nitrogen (AN) and nitrogen deposition exerted the strongest influence, with additional contributions from DBH, slope, and soil sand content. SHAP dependence analyses revealed nonlinear responses across both forest types. In natural forests, FD, CWM SLA, soil sand content, SR, and nitrogen deposition exhibited a positive relationship with productivity (Figure 4a). In planted forests, productivity was positively associated with soil AN, nitrogen deposition, DBH, and soil sand content (Figure 4b). Slope exhibited a unimodal relationship with productivity in both forest types.

**FIGURE 3.**
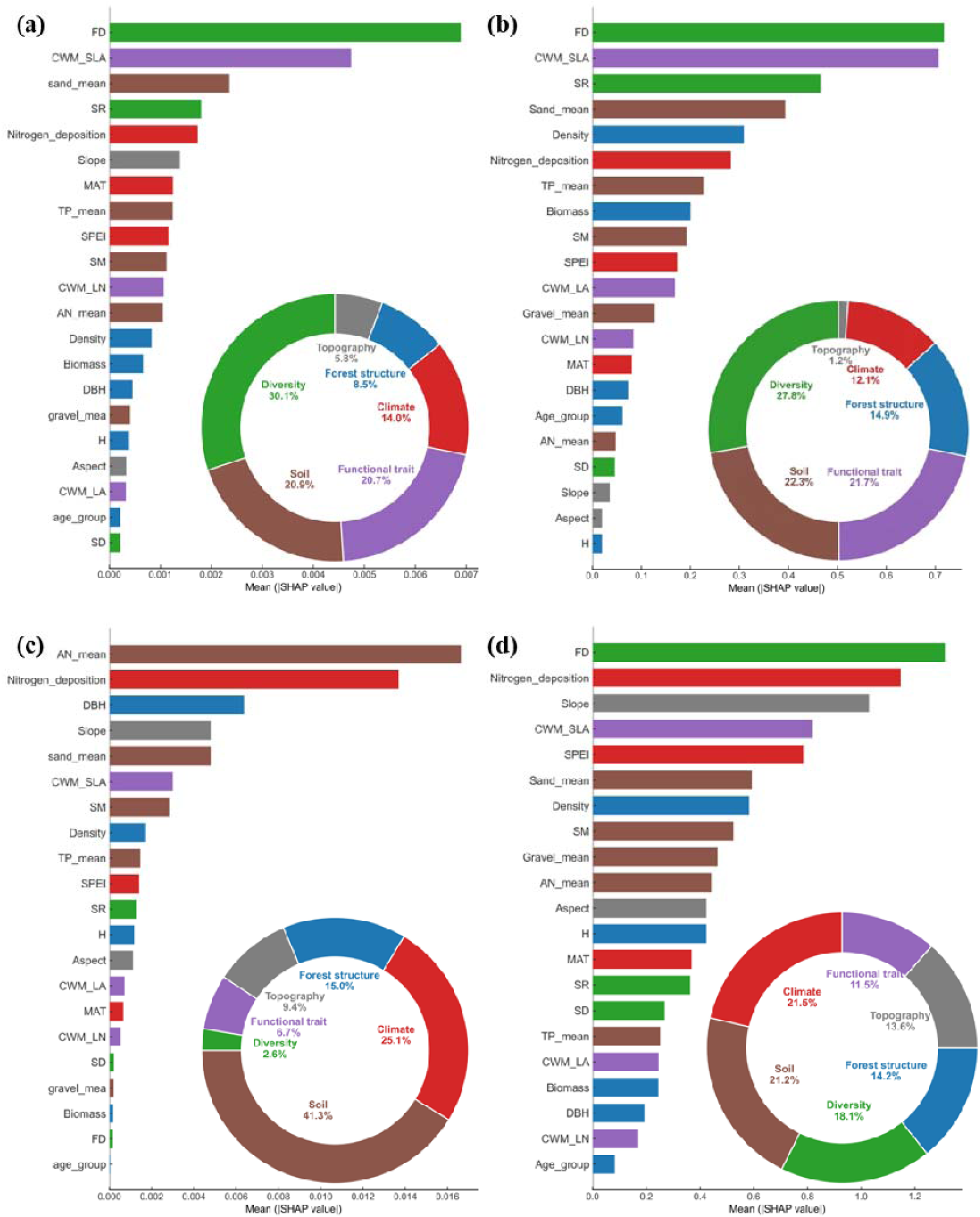
Relative importance of environmental, biological, and structural variables in driving forest productivity and stability across natural and planted forests in China, quantified using mean absolute SHAP values. Results are presented for (a) natural forest productivity, (b) natural forest stability, (c) planted forest productivity and (d) planted forest stability. Horizontal bar charts display the mean |SHAP| value for each predictor variable, reflecting its overall contribution to model predictions, while embedded donut charts illustrate the proportional contribution of six major variable categories: diversity (green), functional trait (purple), forest structure (blue), climate (red), soil (brown), and topography (gray). Variable abbreviations are a follows: Sand_mean, soil sand content; Gravel_mean, soil gravel content; AN_mean, soil alkali-hydrolysable nitrogen; TP_mean, soil total phosphorus; SM, Soil moisture.

**FIGURE 4.**
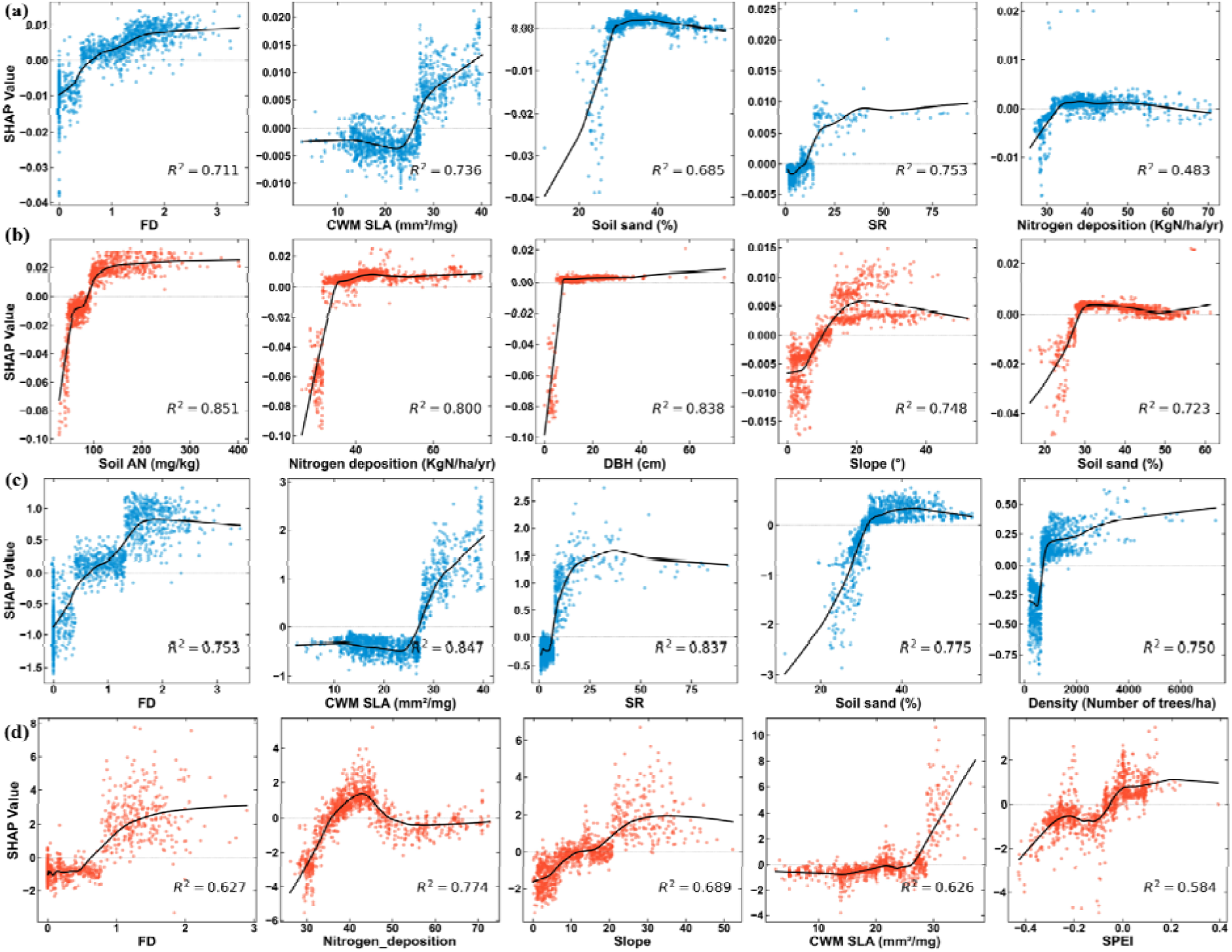
SHAP dependence plots illustrating the nonlinear relationships between the top five most important predictor variables and model output for (a) natural forest productivity, (b) planted forest productivity, (c) natural forest stability and (d) planted forest stability. Blue and orange points represent natural forests and planted forests, respectively, with the fitted smooth curve (black line) representing the marginal effect of each predictor on the SHAP value. Positive SHAP values indicate a positive contribution to the predicted outcome, while negative SHAP values indicate a suppressing effect. The coefficient of determination (R²) is reported for each relationship, indicating the goodness of fit of the smoothed trend.

The relative importance of predictor categories for stability largely mirrored those observed for productivity in natural forests, with tree diversity (27.8%), soil properties (22.3%), and functional traits (21.7%) accounting for a large proportion of the total contribution. The most influential individual predictors were FD, CWM SLA, SR, soil sand content, and stand density (Figure 3b). In planted forests, stability was dominated by climate (21.5%), soil properties (21.2%), and tree diversity (18.1%), with FD, nitrogen deposition, CWM SLA, and the SPEI showing higher contributions (Figure 3d). SHAP dependence results reflected that increased FD, CWM SLA, SR, soil sand content and density were associated with higher stability in natural forests (Figure 4c). In planted forests (Figure 4d), FD, slope, CWM SLA, and SPEI showed a positive correlation with stability increased. Nitrogen deposition exhibited a unimodal relationship with stability, with stability increasing at low deposition levels but declining under high deposition rates.

### 3.3 Causal effects of key driving factors on forest productivity and stability in natural forest and planted forest

To further disentangle the causal effects of dominant drivers, SCM was applied to quantify the influence of key predictors on forest productivity and stability (Figure 5). In natural forests, forest productivity increases with rising FD, CWM SLA, SR and nitrogen deposition (Figure 5a). However, the positive effects of SR and nitrogen deposition weakened at higher levels, indicating saturating relationships. Similarly, productivity exhibited a unimodal response to slope, peaking at intermediate slope values and declining under steeper conditions.

**FIGURE 5.**
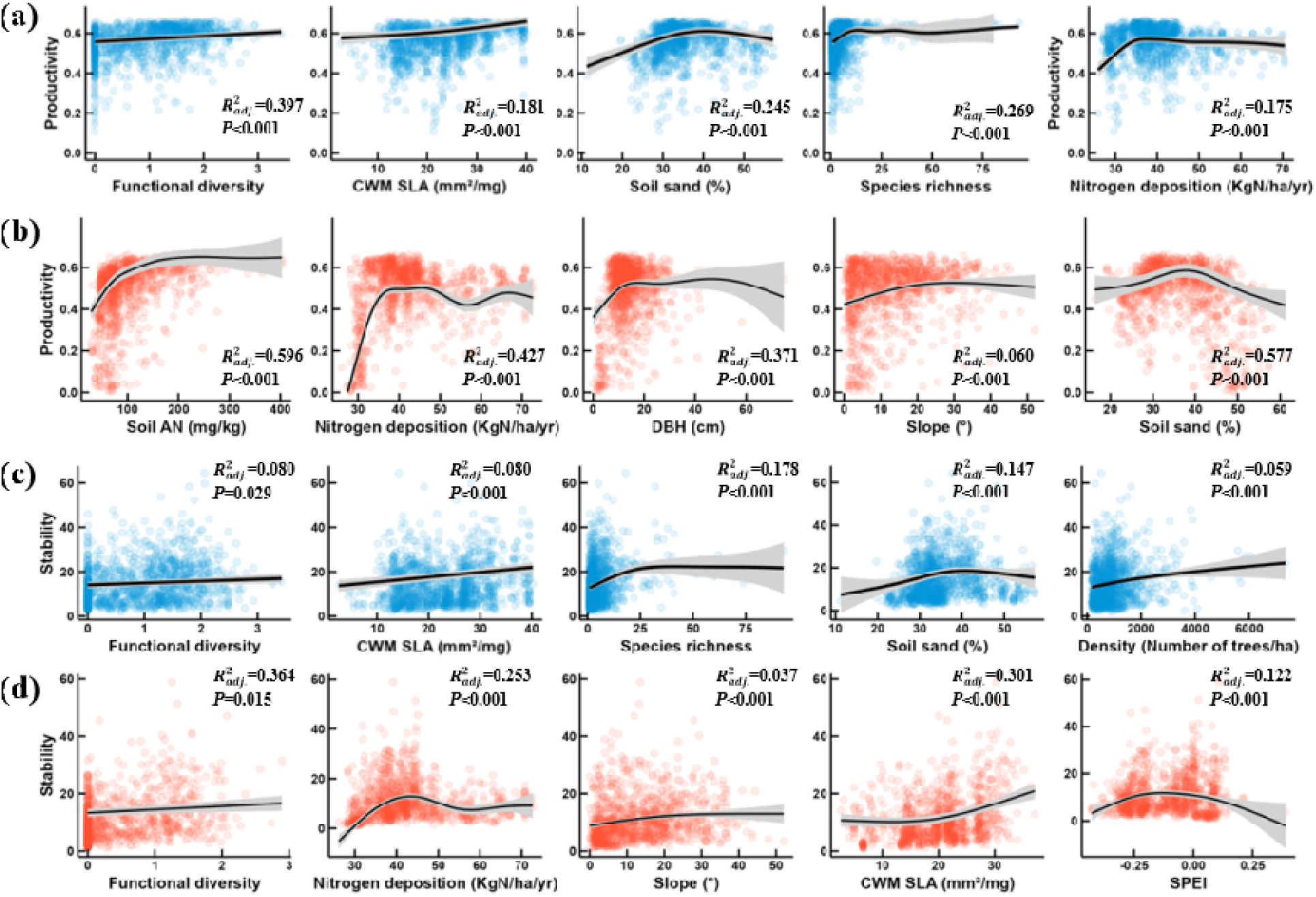
GAMM marginal effect plots illustrating the estimated causal effects of the five most influential predictor variables on (a) natural forest productivity, (b) planted forest productivity, (c) natural forest stability, and (d) planted forest stability. Blue and orange points represent natural forest and planted forest plots, respectively. Shaded ribbons indicate 95% confidence intervals around the fitted smooth functions. The coefficient of determination (R²) and associated p-values are provided for each fitted relationship to evaluate explanatory power and statistical significance.

In planted forests, increasing soil AN and nitrogen deposition exerted strong positive effects on productivity, but these effects plateaued beyond critical thresholds, revealing diminishing returns under nutrient-enriched conditions (Figure 5b). DBH further modulated productivity responses, with productivity increasing at low DBH but gradually declining at higher DBH levels.

In addition, slope and soil sand content exerted unimodal effects on productivity, suggesting optimal productivity under intermediate environmental conditions. Stability increased with FD, CWM SLA, SR, and stand density in natural forests, with the stabilizing effect of SR plateauing in highly diverse forests (Figure 5c). Soil sand content constrained stability through a strong unimodal pathway, with stability peaking at intermediat levels before declining. In planted forests, FD maintained a positive linear effect on stability, whereas CWM SLA showed a nonlinear positive relationship (Figure 5d). In contrast, nitrogen deposition, slope, and SPEI all showed hump-shaped relationships, indicating that stability maximized at intermediate levels of these drivers and declining beyond their respective optima.

## 4. Discussion

We compiled a nationwide forest dataset based on 2,389 plots to compare productivity and stability between natural and planted forests across China and to identify the key drivers and underlying mechanisms from 38 biotic, structural, climatic, and edaphic variables. We hypothesized that natural forests maintain higher productivity and stability (hypothesis 1) and are primarily regulated by biotic attributes, especially tree diversity and stand structure, whereas abiotic factors have stronger effects in planted forests (hypothesis 2). To test these hypotheses, we developed a novel analytical framework combining machine learning (XGBoost), model interpretation (SHAP), and causal inference (SCM). The XGBoost–SHAP approach provides a powerful tool for detecting key predictors and characterizing complex nonlinear relationships between biotic and abiotic factors and forest productivity and stability. On the other hand, the SCM enables explicit causal inference by accounting for confounding structures and spatial dependence. Their combination allows for a more robust and mechanistic understanding of the drivers of productivity and stability in natural and planted forests.

### 4.1. Higher forest productivity and stability in natural forests than in planted forests across China

Our results demonstrate that natural forests exhibit significantly higher productivity and stability than planted forests across China, supporting our first hypothesis. Although previous studies have shown that natural forests are generally more productive than planted forests, our study extends these findings by simultaneously evaluating productivity and stability at the national scale. More importantly, our results further suggest that these advantages are closely associated with the substantial differences in forest structure and tree diversity between natural and planted forests. Compared with planted forests, natural forests in our dataset exhibited larger tree size, higher biomass, species richness, functional diversity, and structural diversity (Figure S2). Larger trees contribute disproportionately to stand-level productivity through their extensive crowns, deeper rooting systems, and greater access to resources (Brown et al. 2004; Collalti et al. 2020; Bordin et al. 2021). Large trees also buffer ecosystem responses to environmental fluctuations and disturbances, thereby enhancing stability (Stephenson et al. 2014; Lutz et al. 2018). Meanwhile, the greater tree diversity and structural complexity observed in natural forests generate complementary resource-use strategies and niche partitioning (Williams et al. 2017; Deng et al. 2025). Vertically stratified canopies enhance light capture, while complementary rooting strategies improve access to soil water and nutrients (Sun et al. 2017; Deng et al. 2025). Such complementary resource use increases both productivity and temporal stability, consistent with biodiversity–ecosystem functioning theory (Tilman 1999). In contrast, planted forests are often characterized by simplified structure, lower tree diversity, and more synchronized growth dynamics, making them more vulnerable to climatic extremes and biotic disturbances such as drought, pest outbreaks, and diseases (Gadgil & Bain 1999; Zhong et al. 2021; Ma et al. 2025).

Nevertheless, the magnitude of these differences varied across ecoregions and stand development stages. Natural forests exhibited stronger advantages in temperate and plateau regions, where have benefited from long-term implementation of the National Natural Forest Conservation Program and restoration efforts in ecologically fragile areas (Zhang et al. 2000). Notably, planted forests in the mid-temperate coniferous region displayed higher stability than natural forests. Two non-exclusive mechanisms may explain this pattern. First, plantations in this region are commonly dominated by drought- and cold-tolerant conifer species (e.g., *Larix gmelinii* and *Pinus sylvestris* var. *mongolica*) that were selectively established to cope with harsh climatic conditions and thus exhibit relatively stable growth under climate variability (Guo et al. 2022; Li et al. 2023). Second, natural forests in this region frequently occur in ecological transition zones and are more exposed to climate warming and intensified drought stress, which can destabilize growth dynamics (Li et al. 2025a). We also found that differences between forest types gradually diminished with stand age. As forests mature, productivity generally declines because of increasing hydraulic constraints, nutrient limitation, and maintenance costs (Ryan et al. 2006; Couvreur et al. 2018; West 2024; Shan et al. 2025). Old-growth natural forests may become increasingly sensitive to climatic extremes such as prolonged drought or heat stress (Bennett et al. 2015), leading to reduced stability. Meanwhile, planted forests tend to develop more complex structure and richer understory communities (Bauhus et al. 2009; Grieco et al. 2024), partially compensating for their initially simplified conditions. These converging trajectories likely explain the reduced differences observed in older stands.

### 4.2 Contrasting driving mechanisms of forest productivity and stability between natural forests and planted forests

Consistent with our second hypothesis, our results reveal a shift from biotic regulation in natural forests to abiotic regulation in planted forests. In natural forests, productivity was primarily regulated by FD, SR, and CWM SLA. These findings reinforce previous evidence that diversity constitutes a stronger driver of productivity than climatic or topographic factors (Hooper et al. 2012; Duffy et al. 2017) and highlight the central role of diversity–productivity relationships in natural forest ecosystems. The positive relationship between FD and productivity provides strong support for the niche complementarity hypothesis, indicating that communities composed of species with contrasting resource-use strategies can exploit resources more efficiently through spatial, temporal, and functional niche partitioning (Williams et al. 2017; Deng et al. 2025). SR also positively influenced productivity, although its effect saturated at higher diversity levels, suggesting increasing functional redundancy once most functional niches become occupied (Loreau et al. 2001; Loreau 2004). Simultaneously, the positive effect of CWM SLA supports the mass-ratio hypothesis (Grime 1998), indicating that dominant species traits play a critical role in determining ecosystem functioning and communities dominated by species with acquisitive resource-use strategies are capable of higher carbon assimilation rates and greater productivity (Wright et al. 2004). In contrast, biomass did not emerge as a dominant explanatory variable per se, suggesting that the vegetation quantity hypothesis plays a less prominent (but not null) role compared to the niche complementarity and mass ratio hypotheses. Together, our study provides new empirical evidence for the relative contributions of these mechanisms and suggests that productivity in natural forests is jointly regulated by niche complementarity and mass-ratio effects. In contrast to natural forests, productivity in planted forests was regulated primarily by abiotic factors, including climate, soil conditions, and nitrogen deposition. This shift likely arises from the inherently low species and functional diversity of plantations, which constrains niche differentiation and diminishes biotic feedbacks between vegetation and soil (Liao et al. 2012). Simplified stand structures and homogenized litter inputs may further reduce the complexity of soil microbial networks (Qu et al. 2024), thereby weakening internal ecosystem regulation and increasing dependence on external resource availability. Among all predictors, soil AN and nitrogen deposition emerged as the strongest determinants of planted forest productivity. Nitrogen is widely recognized as a key limiting nutrient in terrestrial ecosystems (LeBauer & Treseder 2008; Li et al. 2025b), and our analyses revealed clear nonlinear responses consistent with nitrogen saturation theory. Productivity increased with nitrogen availability under low to moderate inputs but reached a plateau or declined when nitrogen supply became excessive. Moderate nitrogen inputs likely stimulate plant growth and microbial nutrient cycling (O’Sullivan et al. 2019; Luo et al. 2022), whereas excessive nitrogen deposition can induce soil acidification, disrupt microbial communities, and reduce biological diversity, ultimately constraining productivity (Bobbink et al. 2010; Tian & Niu 2015; Zhou et al. 2020). Similar nonlinear responses were observed for topographic and edaphic variables, including slope and soil sand content, highlighting the existence of environmental optima beyond which productivity declines.

Patterns of stability partly mirrored those observed for productivity but revealed important mechanistic differences. Whereas productivity in planted forests became increasingly dependent on abiotic conditions, stability remained strongly associated with tree diversity across both forest types. Functional diversity emerged as the most influential predictor of stability in both natural and planted forests, while species richness and functional traits also contributed substantially, particularly in natural forests. The positive effects of functional diversity and species richness are consistent with niche complementarity and insurance effects (Yachi & Loreau 1999). Forest communities with higher tree diversity are more likely to contain species that differ in resource-use strategies, phenology, and stress tolerance (Loreau & De Mazancourt 2013). Such differences generate asynchronous responses to environmental variability, allowing declines in some species to be compensated by increases in others and thereby stabilizing ecosystem functioning through time (Loreau & De Mazancourt 2013). Interestingly, communities characterized by higher CWM SLA exhibited greater stability in both natural and planted forests. Although previous studies often emphasize the stabilizing role of conservative resource-use strategies (Craven et al. 2018; Valerio et al. 2022; Luo et al. 2023), our results suggest that acquisitive species may enhance stability through rapid recovery following disturbance (Bazzichetto et al. 2024; Cui et al. 2026). Species with high SLA generally possess greater photosynthetic capacity and faster growth rates, thereby enhancing stability at both the species and community levels (Huang et al. 2022). Despite the importance of tree diversity, abiotic factors exerted stronger influences on stability in planted forests. Nitrogen deposition displayed a hump-shaped relationship with stability in planted forests, likely mediated through biodiversity pathways. Previous research has demonstrated a unimodal relationship between nitrogen deposition and plant diversity, whereby moderate nitrogen inputs enhance diversity but excessive deposition leads to biodiversity loss (Simkin et al. 2016), ultimately reducing stability. Moreover, the stronger response of planted forests to SPEI indicates that planted forests are more vulnerable to hydroclimatic variability than natural forests, consistent with previous evidence that plantation ecosystems often exhibit greater drought sensitivity (Zhong et al. 2021; Ma et al. 2025).

### 4.3 Implications for future afforestation and forest management

Our findings have important implications for future afforestation and forest management. Over recent decades, large-scale afforestation programmes have substantially expanded forest area across China, but many newly established forests remain dominated by monoculture plantations. Our results indicate that improving forest productivity and stability requires moving beyond a sole focus on forest area expansion and placing greater emphasis on the quality and diversity of forest communities. In particular, functional diversity emerged as a stronger and more consistent predictor of both productivity and stability than species richness alone. Future afforestation and forest management efforts should therefore prioritize the assembly of species with complementary functional traits rather than simply increasing the number of species present. Our findings also suggest the need to incorporate resource-acquisition or fast-growing species, such as those with higher SLA, to enhance both productivity and post-disturbance recovery capacity. In addition to tree diversity, our results highlight the importance of nitrogen management and climate adaptation strategies. While moderate nitrogen availability can enhance forest productivity, excessive nitrogen deposition may undermine both productivity and stability. Forest management should therefore balance nutrient inputs while minimizing the ecological risks associated with nitrogen enrichment. At the same time, the greater drought sensitivity of planted forests suggests an urgent need to improve plantation resilience under future climate change. Increasing functional diversity, selecting drought-tolerant species, and promoting structurally complex forest communities may represent effective strategies for enhancing ecosystem resistance and recovery.

## Data availability

The raw data that support the findings of this study are publicly available on the Figshare Repository at: https://doi.org/10.6084/m9.figshare.32956802.

## Code availability

The code that supports the findings of this study is publicly available on the Figshare Repository at: https://doi.org/10.6084/m9.figshare.32956802.

## Author Contributions

**Mingjie Chen**: conceptualization, writing – original draft, investigation, methodology, visualization, writing – review and editing, formal analysis. contributed to conceiving and designing the study, conducting the data analysis, developing the methodology, writing the original manuscript, and revising the manuscript. **Daoli Peng**: supervision, project administration, writing – review and editing, funding acquisition. **Elia Vangi**: writing – review and editing. **Mauro Morichetti**: writing – review and editing. **Alessio Collalti**: conceptualization, writing – review and editing.

## Acknowledgments

This study was supported by the National Key Research and Development Program of China (grant number 2023YFD2200403). We are grateful to Daniela Dalmonech and Paulina F. Puchi for their insightful comments and constructive suggestions on earlier versions of the manuscript. We would also like to thank the editors and anonymous reviewers for their valuable suggestions in improving the quality of the manuscript.

