## Supplementary material for "Natural and planted forests differ in the mechanisms regulating productivity and stability"

**Table S1** Summary of the environmental variables used in this study.

| Variables | Period | Resolution | Source |
| --- | --- | --- | --- |
| Mean annual temperature and mean annual precipitation | 2001-2020 | 1km | (Peng et al. 2019) |
| Standardized precipitation evapotranspiration index | 2001-2020 | 1km | (He et al. 2025) |
| Nitrogen deposition | 2005-2020 | 0.25° | (Zhou et al. 2023) |
| Atmospheric carbon dioxide concentration | 2015-2020 | 1km | (He et al. 2023) |
| Elevation, aspect and slope | 2001-2020 | 30m | (NASA JPL, 2020) |
| Soil moisture | 2001-2020 | 1km | (Zhang et al. 2023) |
| Soil sand, silt, clay, rock fragment, porosity, bulk density, pH, organic carbon, cation exchange capacity, total nitrogen, total phosphorus, total potassium, alkali-hydrolysable nitrogen, available potassium, Available phosphorous | 1970-2010 | 1km | (Shi et al. 2025) |

**Table S2** Optimal hyperparameters for each model.

| Model | n_estimators | max_depth | learning_rate | min_child_weight |
| --- | --- | --- | --- | --- |
| Natural forest productivity | 200 | 8 | 0.002 | 1 |
| Planted forest productivity | 200 | 6 | 0.003 | 1 |
| Natural forest stability | 200 | 4 | 0.004 | 1 |
| Planted forest stability | 500 | 6 | 0.3 | 1 |

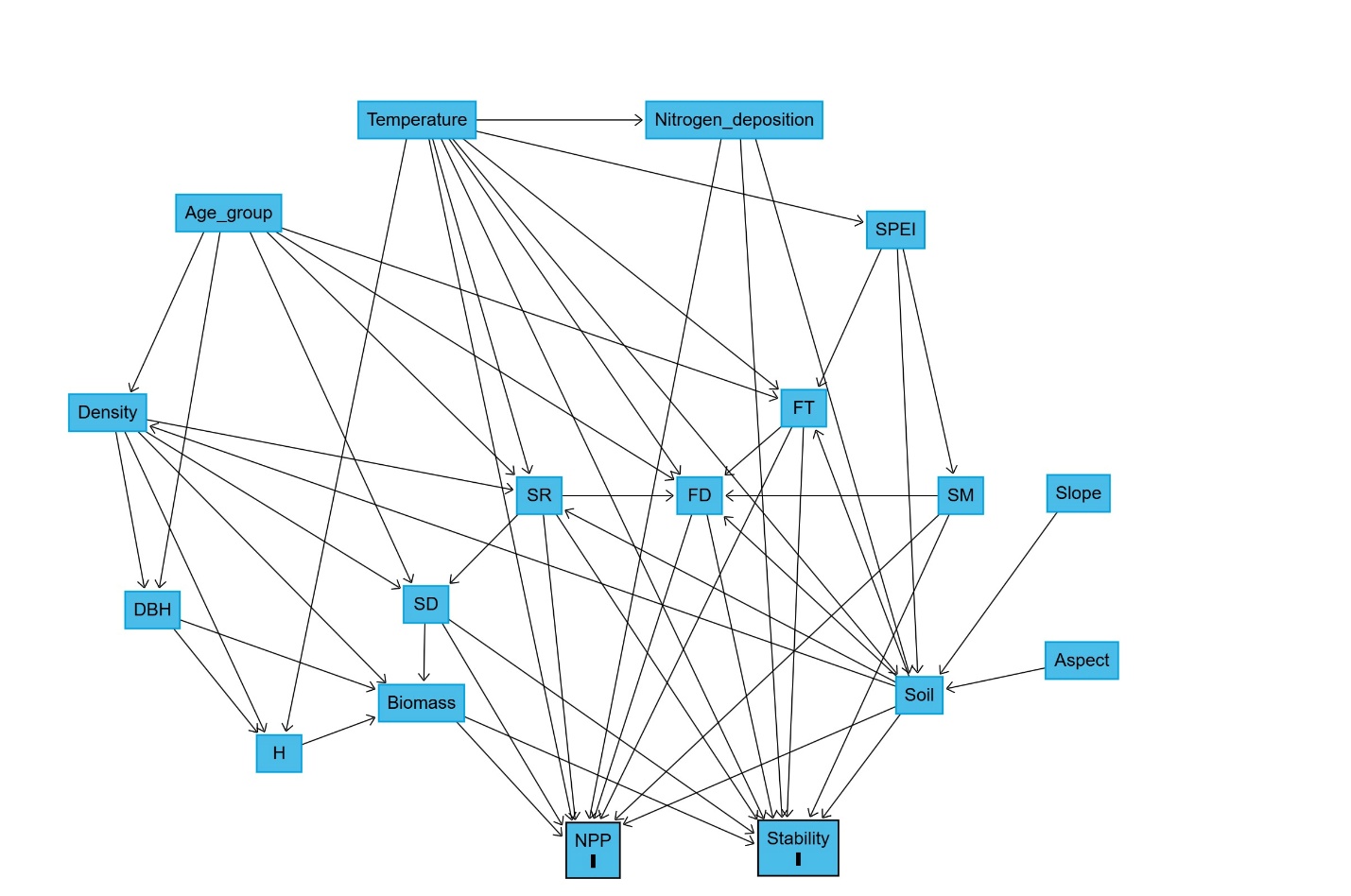

**FIGURE S1** Constructed Directed Acyclic Graph to depict the causal structure of forest productivity and stability. SD, structural diversity; SR, species richness; FD, functional diversity; FT, functional trait; SM, soil moisture.

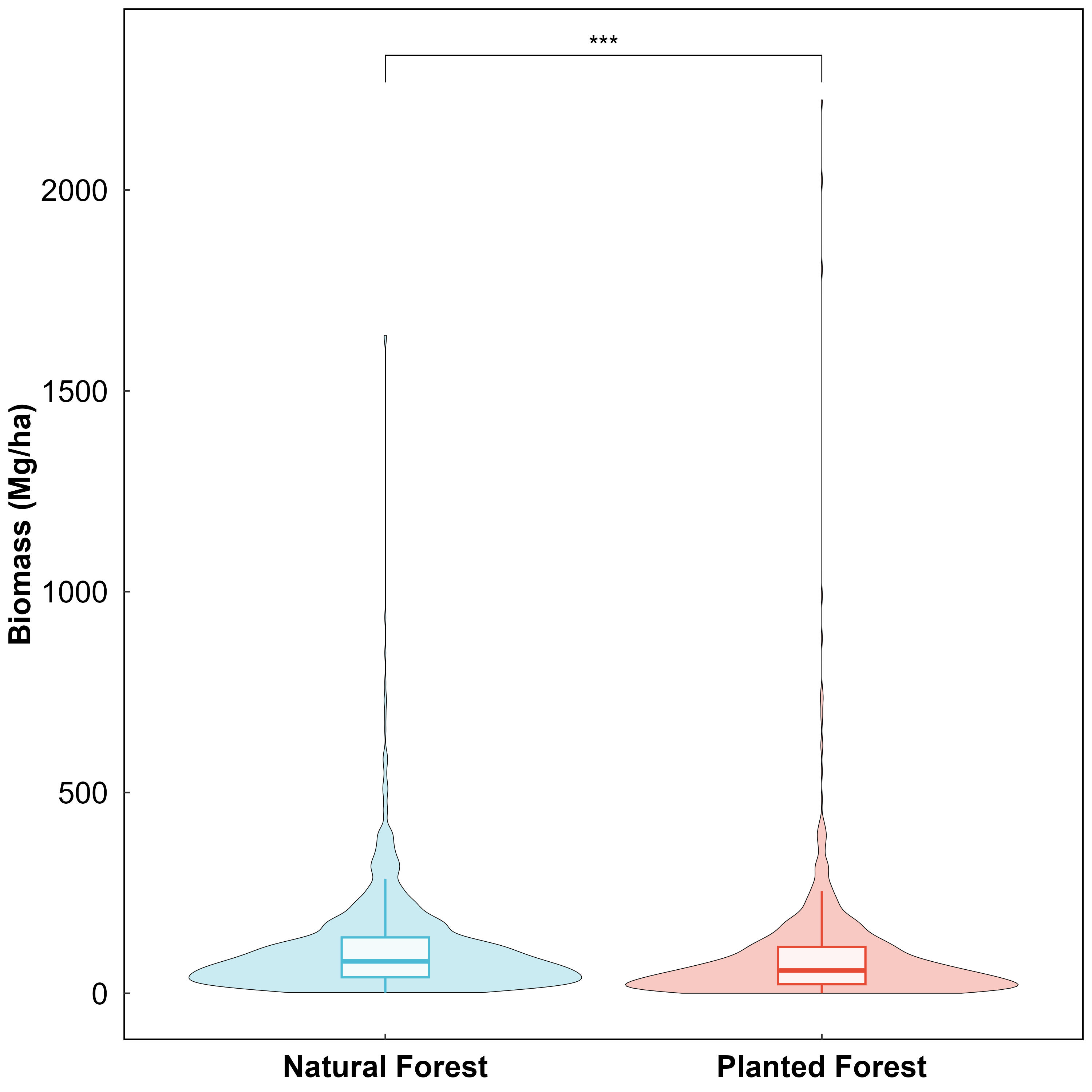

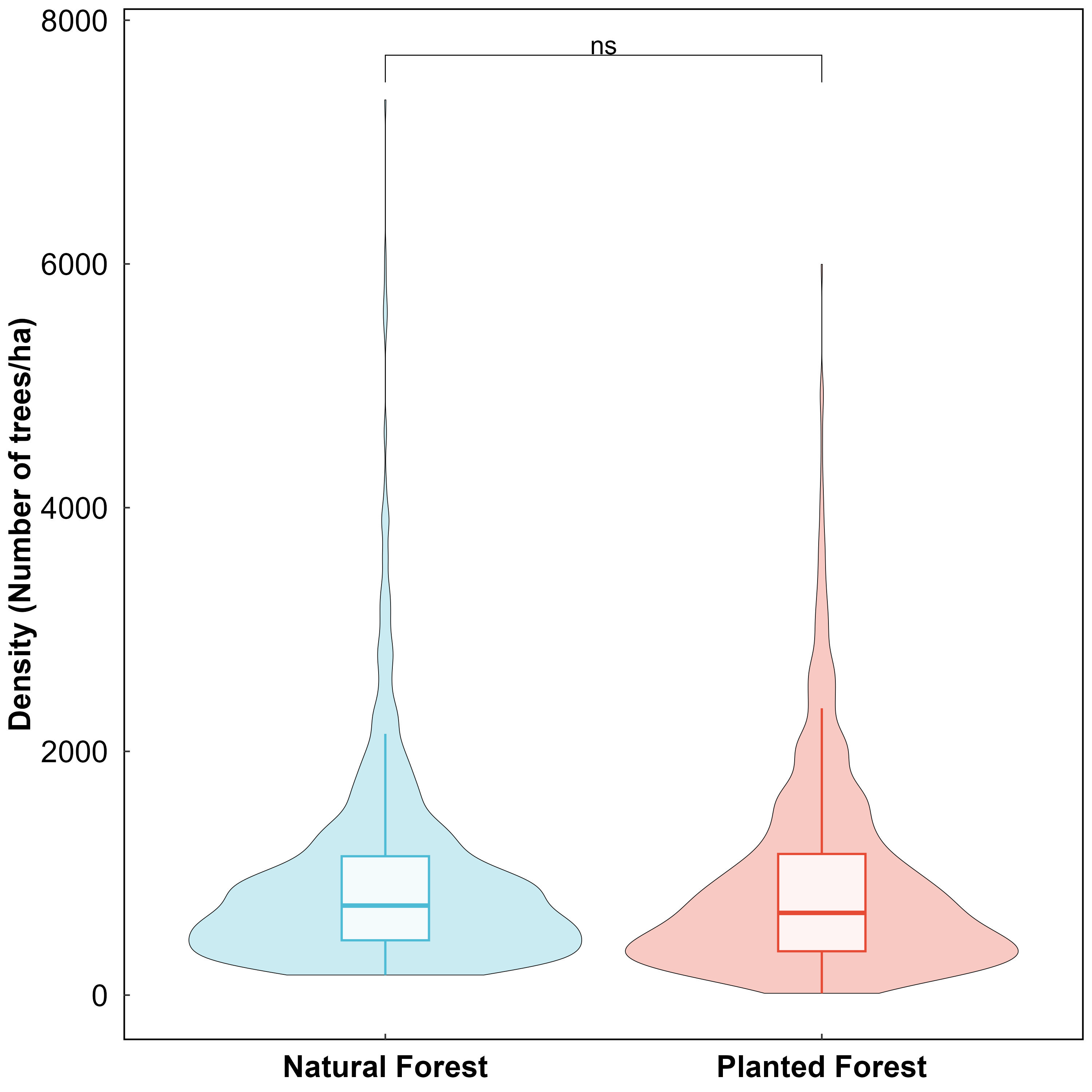

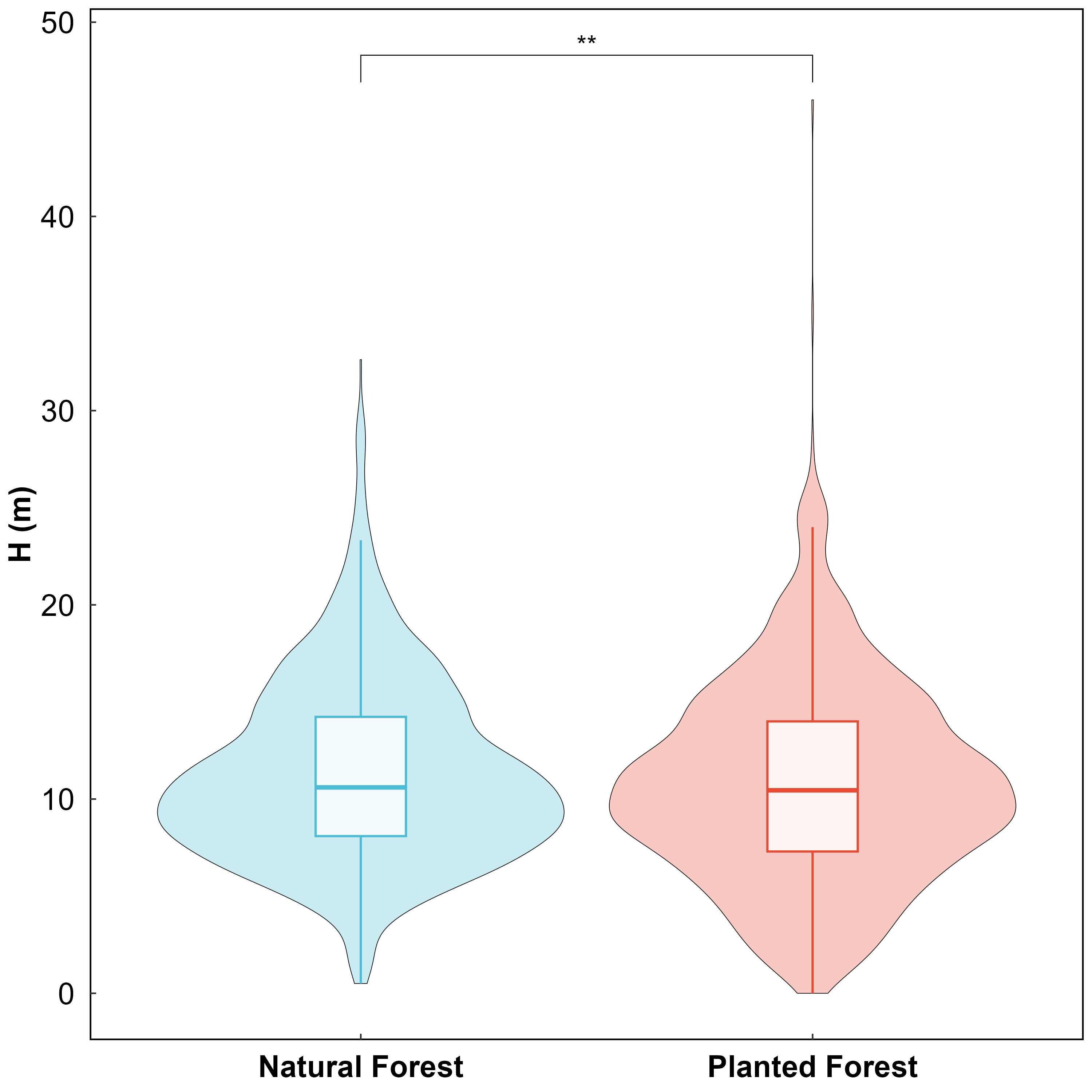

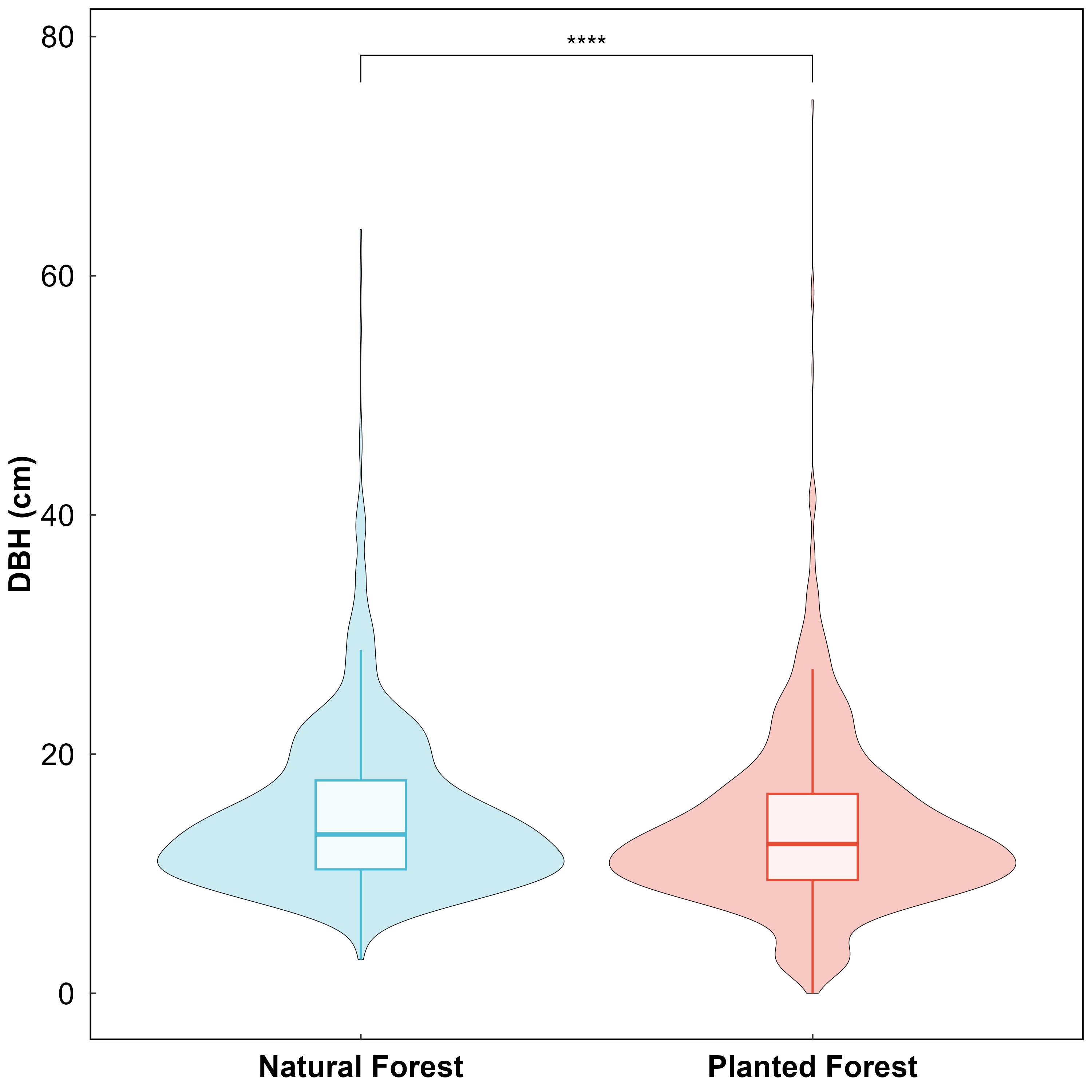

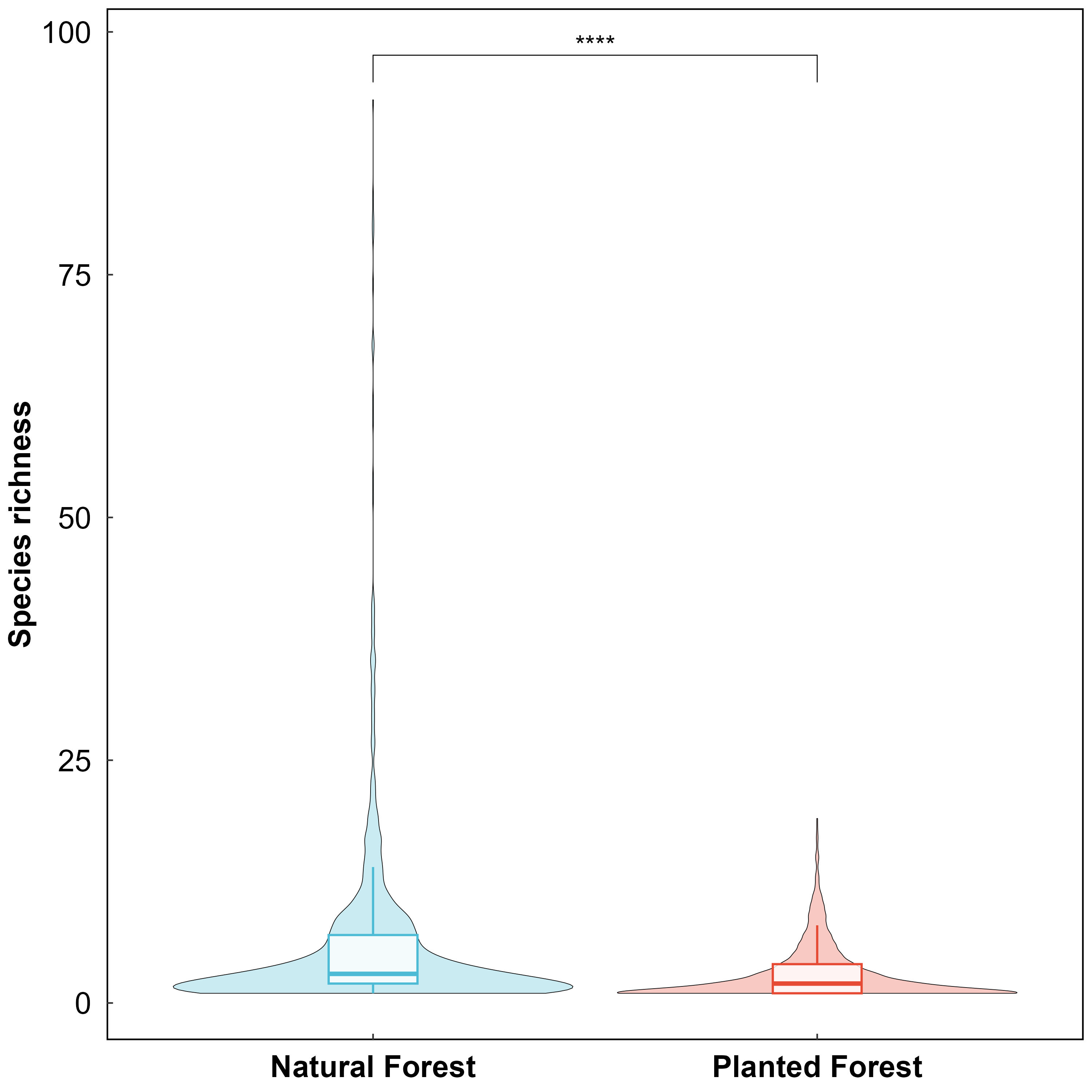

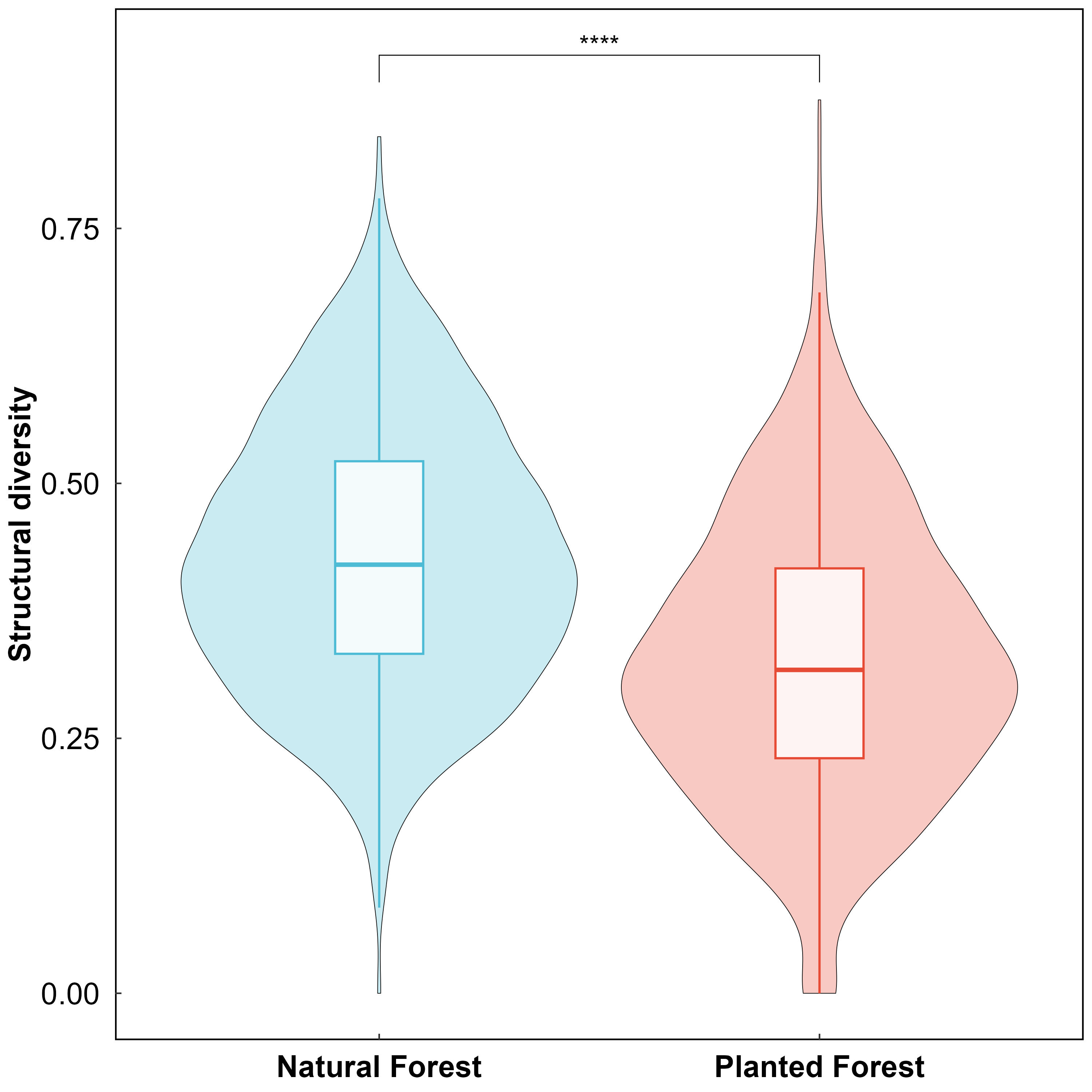

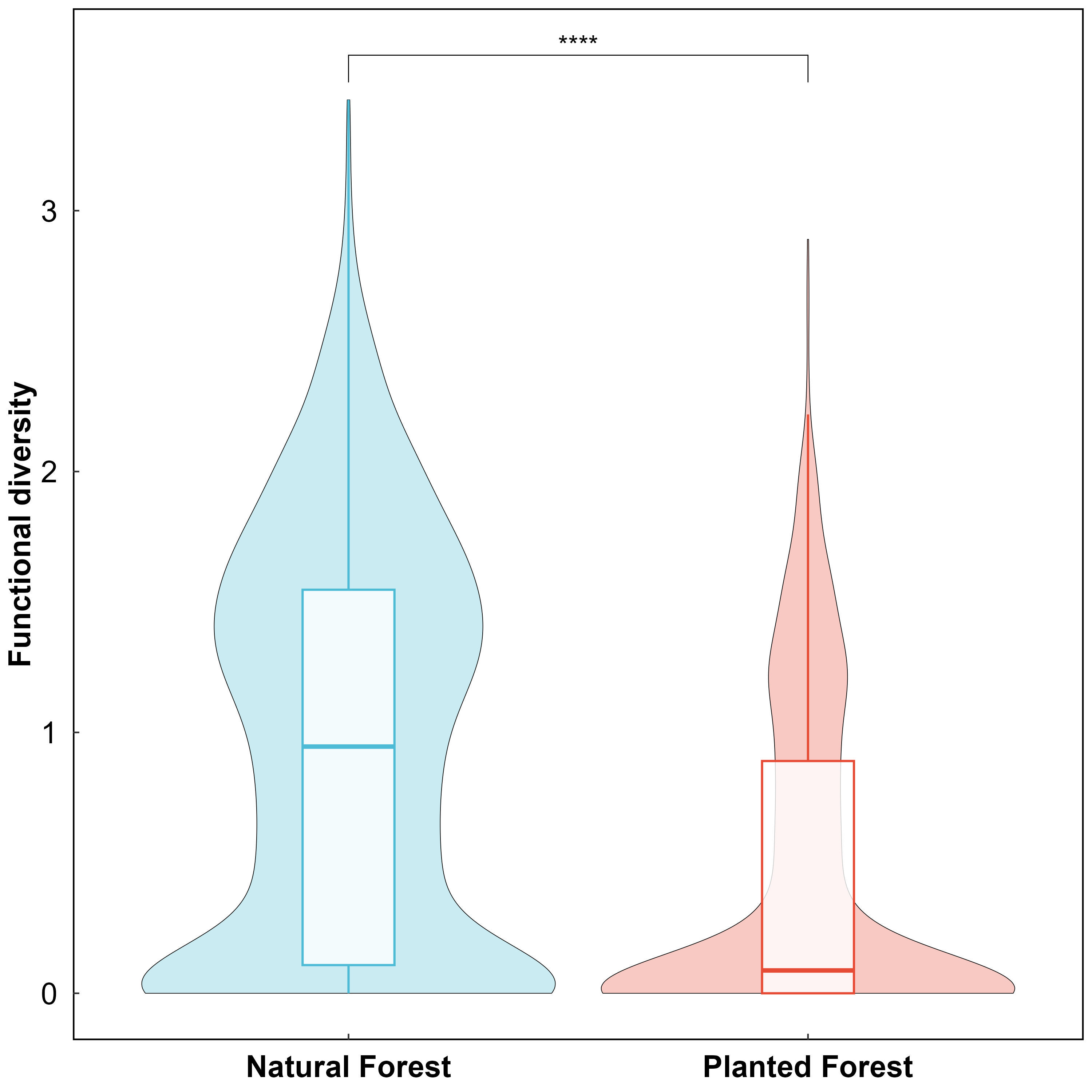

**FIGURE S2** Statistical results on the differences in forest structures between natural and planted forests across China. The symbols indicate the significance levels of the ANOVA: **** for p<0.0001, *** for p<0.001, ** for p<0.01, * for p<0.05, and ns for p>0.05.

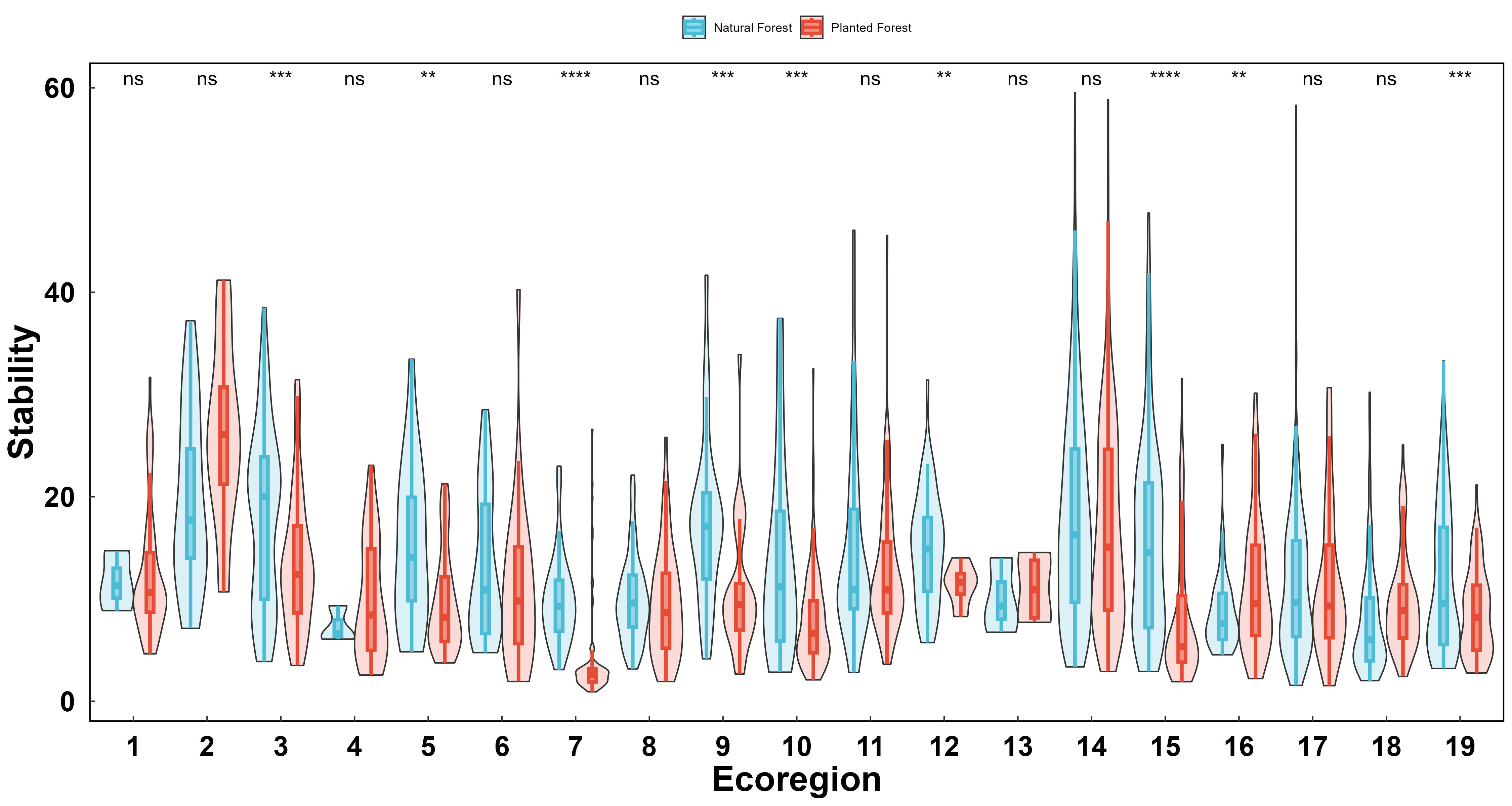

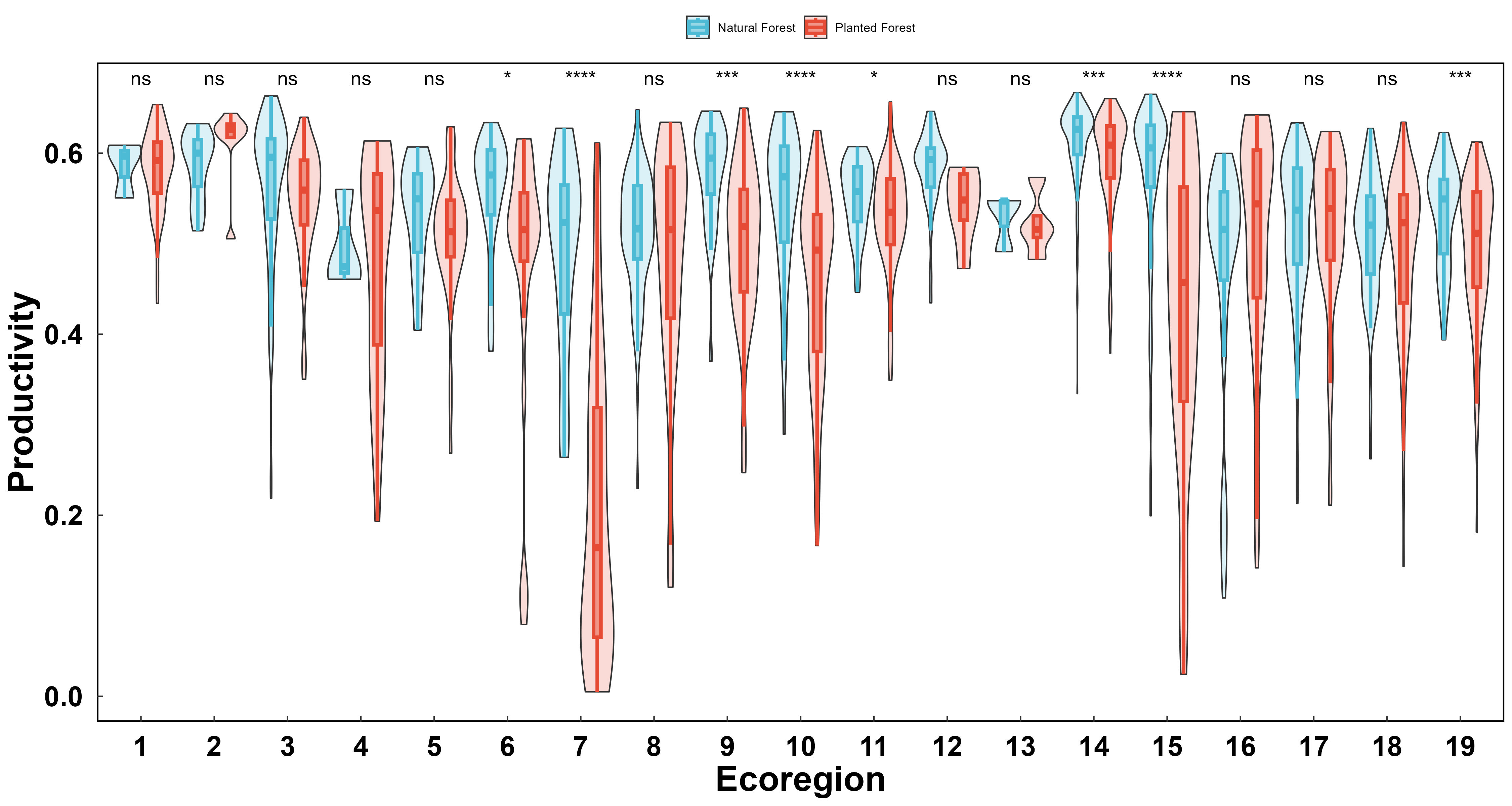

**FIGURE S3** Statistical results on the differences in productivity and stability between natural and planted forests across ecoregions in China. The numbers represent the specific ecoregions: 1: Northern tropical broadleaf forest; 2: Northern temperate mixed forest; 3: Northern subtropical mixed forest; 4: Northern subtropical broadleaf forest; 5: Northern subtropical coniferous forest; 6: Plateau climate mixed forest; 7: Plateau climate broadleaf forest; 8: Plateau climate coniferous forest; 9: Southern temperate mixed forest; 10: Southern temperate broadleaf forest; 11: Southern temperate coniferous forest; 12: Southern subtropical mixed forest; 13: Southern subtropical coniferous forest; 14: Mid-temperate mixed forest; 15: Mid-temperate broadleaf forest; 16: Mid-temperate coniferous forest; 17: Mid-subtropical mixed forest; 18: Mid-subtropical broadleaf forest; 19: Mid-subtropical coniferous forest. The symbols indicate the significance levels of the ANOVA: **** for *p*<0.0001, *** for *p*<0.001, ** for *p*<0.01, * for *p*<0.05, and *ns* for *p*>0.05.

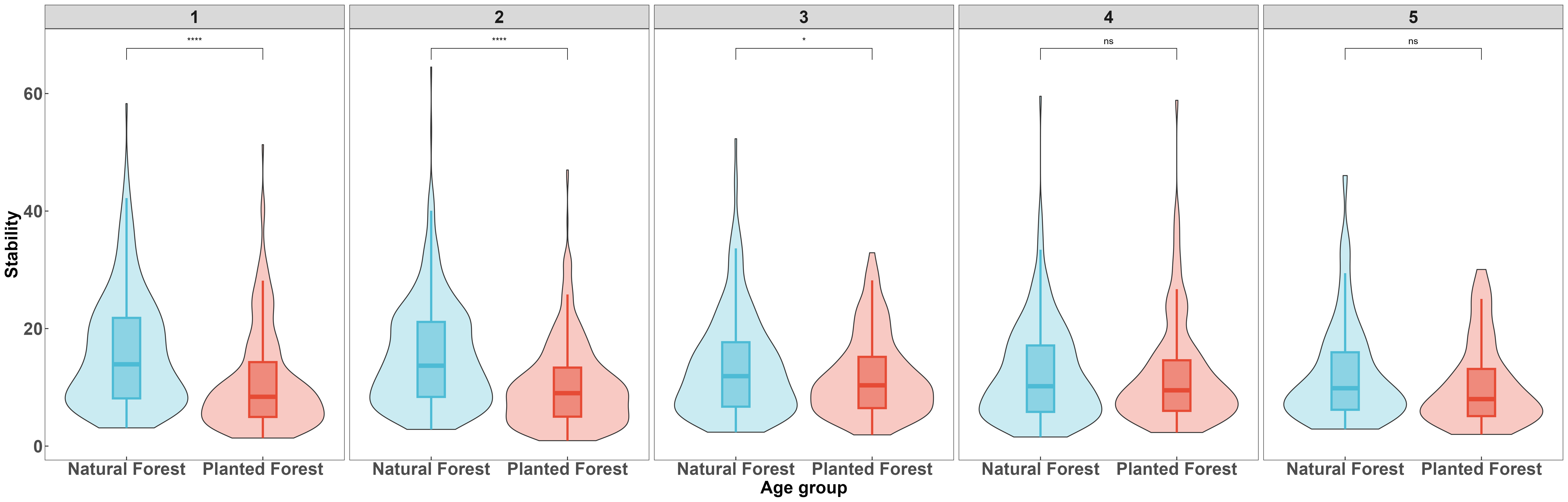

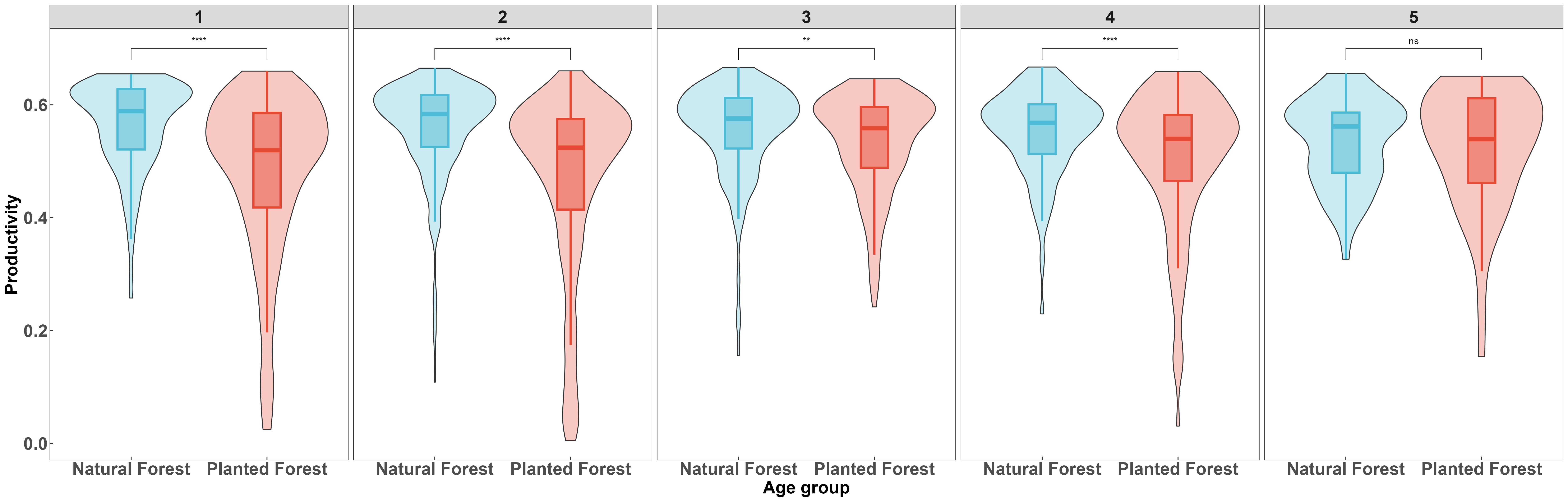

**FIGURE S4** Statistical results on the differences in productivity and stability between natural and planted forests across forest age groups in China. The numbers represent the specific age groups: 1: Young forests; 2: Middle-aged forests; 3: Near-mature forests; 4: Mature forests; 5: Over-mature forests. The symbols indicate the significance levels of the ANOVA: **** for *p*<0.0001, *** for *p*<0.001, ** for *p*<0.01, * for *p*<0.05, and *ns* for *p*>0.05.
